# BRCA2 stabilises RAD51 and DMC1 nucleoprotein filaments through a conserved interaction mode

**DOI:** 10.1101/2023.11.20.567968

**Authors:** James M. Dunce, Owen R. Davies

## Abstract

BRCA2 is essential for DNA repair by homologous recombination in mitosis and meiosis. It interacts with recombinases RAD51 and DMC1 to facilitate the formation of nucleoprotein filaments on resected DNA ends that catalyse recombination-mediated repair. BRCA2’s BRC repeats bind and disrupt RAD51 and DMC1 filaments, whereas its PhePP motifs binds to recombinases in a manner that stabilises their nucleoprotein filaments. However, the mechanism of filament stabilisation has hitherto remained unknown. Here, we report the crystal structure of a BRCA2-DMC1 complex, revealing how PhePP motifs bind to recombinases. This novel mode of interaction is conserved for RAD51 and DMC1, which selectively bind to BRCA2’s two distinct PhePP motifs. In both cases, BRCA2 PhePP motifs enhance the stability of nucleoprotein filaments, protecting them from BRC-mediated disruption. Hence, we report the structural basis of how BRCA2’s PhePP motifs stabilise RAD51 and DMC1 nucleoprotein filaments for their essential roles in mitotic and meiotic recombination.

## Introduction

DNA double-strand break (DSB) repair by homologous recombination is critical for genome integrity and fertility (Page and Hawley, 2003, Wright et al., 2018, Sun et al., 2020). In somatic cells, DSBs arise due to exogenous damage and upon replication fork collapse (Wright et al., 2018). These lesions can be repaired, and replication restarted, through inter-sister recombination, of which the machinery has an additional role in protecting stalled replication forks (Feng and Jasin, 2017, Tye et al., 2021). In meiosis, a programme of DSB induction triggers inter-homologue recombination (Page and Hawley, 2003), enabling synapsis between homologues (Adams and Davies, 2023), and the formation of crossovers that ensure correct chromosome segregation and enhance genetic diversity (Hunter, 2015, Zickler and Kleckner, 2015). Hence, defects in homologous recombination are associated with chromosome instability, increased cancer risk and infertility (Hall et al., 1990, Patel et al., 1998, Sharan et al., 2004).

The tumour suppressor BRCA2 performs a central role in the mechanics of recombination by loading recombinases onto resected DNA ends to form nucleoprotein filaments that mediate strand invasion and homology search within the template DNA (Jensen et al., 2010, Liu et al., 2010, Thorslund et al., 2010). This involves the displacement of RPA from newly resected DNA ends (Shahid et al., 2014, Bell et al., 2023), and the remodelling of recombinases from their oligomeric assemblies into protein-DNA filaments that are active in recombination (Jensen et al., 2010, Liu et al., 2010, Thorslund et al., 2010). Whilst RAD51 is the universal recombinase, meiosis also requires DMC1 (Yoshida et al., 1998, Pittman et al., 1998), which enables mismatch-tolerant inter-homologue recombination (Lee et al., 2015). BRCA2 and RAD51 knockouts are embryonically lethal (Tsuzuki et al., 1996, Sharan et al., 1997), their disruption in cell lines leads to gross chromosomal rearrangements (Sonoda et al., 1998, Patel et al., 1998, Yu et al., 2000), and human BRCA2 mutations are strongly associated with early-onset breast and ovarian cancers (Hall et al., 1990). Further, DMC1 disruption, and germline-specific deficiencies of BRCA2 and RAD51, lead to meiotic impairment and infertility (Yoshida et al., 1998, Pittman et al., 1998, Sharan et al., 2004, Dai et al., 2017).

BRCA2 interacts with RAD51 and DMC1 recombinases through BRC repeats and PhePP motifs (Figure 1a) (Sharan et al., 1997, Wong et al., 1997, Davies et al., 2001, Esashi et al., 2005, Thorslund and West, 2007). The eight BRC repeats, located within BRCA2’s central exon 11 region, bind to RAD51 and DMC1 with various affinities (Wong et al., 1997, Carreira and Kowalczykowski, 2011, Martinez et al., 2016). They interact via molecular mimicry with the self-association interface that mediates the oligomeric assembly and nucleoprotein filament formation of recombinases, by locking them in inactive monomeric states (Pellegrini et al., 2002). Hence, BRC repeats are thought to remodel RAD51 and DMC1, from their isolated states in large filamentous structures and octameric rings, respectively, to nucleate their assembly on resected DNA ends into active nucleoprotein filaments. BRCA2 has two PhePP motifs, located within exons 14 and 27 (herein referred to as Ex14 and Ex27, respectively; Figure 1a) (Sharan et al., 1997, Esashi et al., 2005, Thorslund et al., 2007). Whilst Ex14 and Ex27 share an eponymous FxPP motif, their sequences are otherwise divergent (Figure 1a), and they demonstrate specificity for DMC1- and RAD51-binding, respectively (Sharan et al., 1997, Esashi et al., 2005, Thorslund et al., 2007). Importantly, Ex27 binds to RAD51 filaments, rather than monomers, and protects nucleoprotein filaments from BRC-mediated disruption (Davies and Pellegrini, 2007, Esashi et al., 2007). However, it remains unknown how PhePP motifs interact with recombinases.

**Figure 1.**
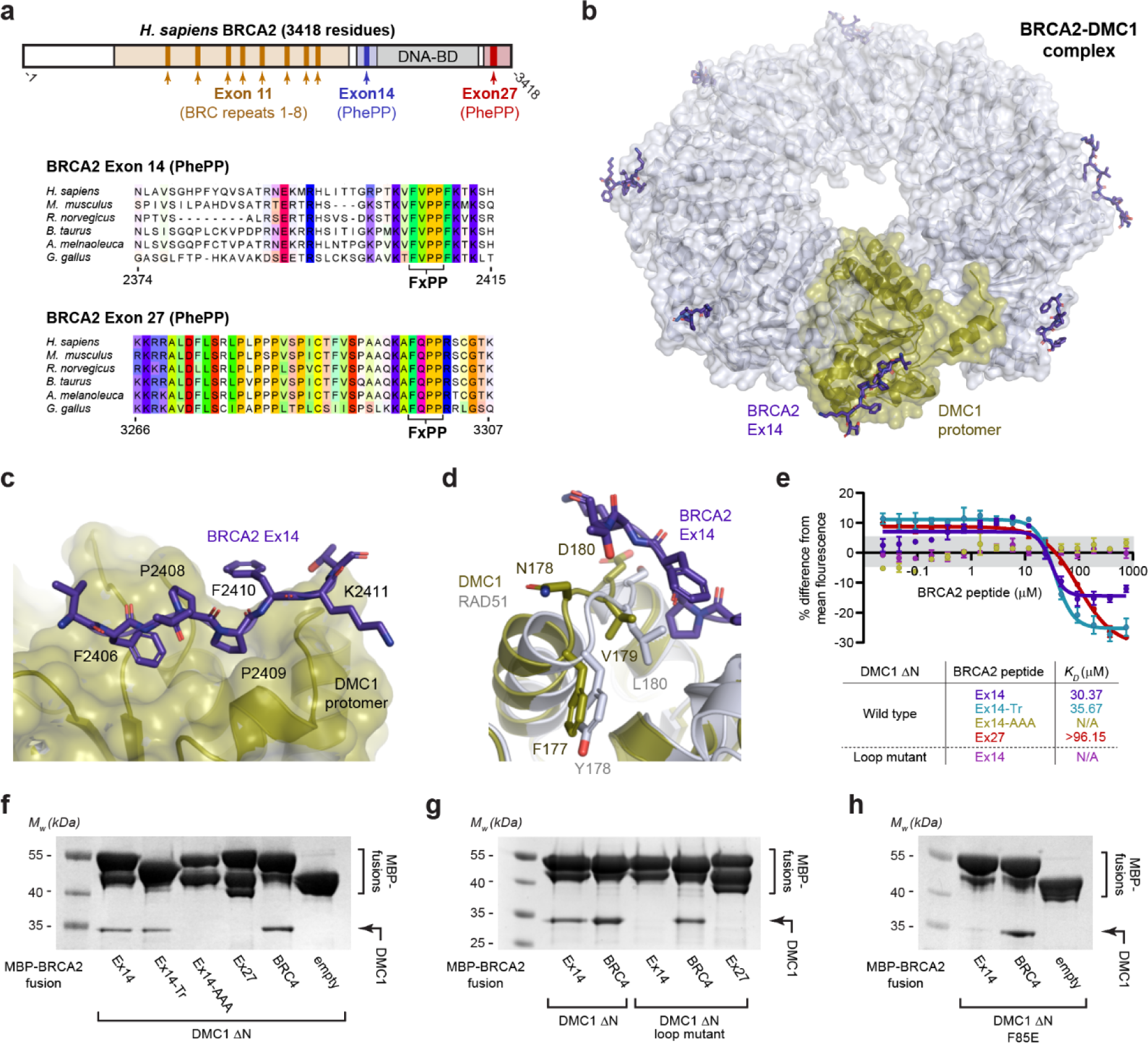
Crystal structure of a BRCA2-DMC1 complex. **(a)** Schematic of the human BRCA2 sequence (top) with multiple sequence alignments of the exon 14 (middle) and exon 27 (bottom) regions, highlighting their PhePP (FxPP) motifs. (**b**) Crystal structure of the complex between BRCA2 Ex14 peptides and a DMC1 ΔN octameric ring. (**c**) Interaction between the PhePP region of the BRCA2-Exon14 peptide (purple) and a DMC1 protomer (yellow), highlighting the FxPP amino-acids F2406, P2408 and P2409. (**d**) Superposition of the BRCA2-DMC1 structure with the RAD51 core, highlighting how the PhePP-binding loop diverges between recombinases, consisting of 177-FNVD-180 in DMC1 and 178-YGLS-181 in RAD51. (**e**) Microscale thermophoresis (MST) of interactions between BRCA2 peptides and DMC1 ΔN. BRCA2 Ex14 and its truncation (for crystallography) bind to DMC1 core with affinities of 30 μM and 36 μM, respectively, whereas interactions are not detectable for the BRCA2-Ex14-AAA mutant (F2406A, P2408A, P2409A), or for the DMC1 ΔN loop mutant (replacing 147-GAGGYPG-153 and 178-NVDHDA-183 with RAD51 amino-acids IDRGGGE and GLSGSD, respectively). BRCA2 Ex27 binds to DMC1 ΔN with an affinity of >96 μM. (**f-h**) Amylose pull-downs of (**f**) DMC1 ΔN, (**g**) DMC1 ΔN wild-type and loop mutant, and (**h**) DMC1 ΔN F85E mutant, following recombinant co-expression with MBP-BRCA2 Ex14 wild-type, truncation (for crystallography) and AAA mutant, Ex27, BRC4 and free MBP (empty).

Here, we report the crystal structure of BRCA2 Ex14 bound to DMC1, and that BRCA2 Ex27 binds to RAD51 through the same core interaction mode. Hence, we uncover the structural basis of how BRCA2 PhePP motifs bind to RAD51 and DMC1 recombinases and stabilise their nucleoprotein filaments for mitotic and meiotic recombination.

## Results

### Crystal structure of a BRCA2-DMC1 complex

The mode of interaction between BRCA2’s PhePP motifs and recombinases has remained unknown since the discovery of this filament-stabilising binding site over two decades ago (Sharan et al., 1997, Esashi et al., 2005, Davies and Pellegrini, 2007, Esashi et al., 2007). As DMC1 exists in octameric rings, whereas RAD51 forms filaments, we reasoned that DMC1 may be more amenable for structure solution of a BRCA2 PhePP-bound complex. Hence, we established a crystallisation system for the structural core of human DMC1 in which its N-terminal domain was deleted (herein referred to as DMC1 ΔN; amino-acids 83-340). We first validated this system by determining the native structure to a resolution of 2.05 Å, using a previous DMC1 structure (PDB accession 4HYY: Du and Luo, 2013) as a template for molecular replacement (Table 1). We then soaked DMC1 ΔN crystals with a peptide corresponding to a truncated form of BRCA2’s Ex14 (herein referred to as Ex14-Tr; amino-acids 2401-2414). Whilst soaking substantially reduced crystal quality, we obtained X-ray diffraction data to anisotropic resolution limits of 3.40–5.80 Å, in which additional peptide density was clearly observed (Figure 1b, Table 1 and Supplementary Figure 1). The peptide density was located adjacent to a loop formed by DMC1 amino-acids 178-183, which is a crystal lattice contact. Hence, the reduction in resolution upon BRCA2 Ex14-binding is owing to the introduction of a defect in the crystal lattice.

**Table 1.** Data collection, phasing and refinement statistics.

|  | DMC1 | BRCA2-DMC1 |
| --- | --- | --- |
| **PDB accession** | 6R3P | 8R2G |
| **Data collection** |  |  |
| Space group | I422 | P4<sub>1</sub>22 |
| Cell dimensions |  |  |
| *a*, *b*, *c* (Å) | 174.49, 174.49, 179.57 | 125.40, 125.40, 364.195 |
| $\alpha$, $\beta$, $\gamma$ (°) | 90, 90, 90 | 90, 90, 90 |
| Resolution (Å) | 47.49 – 2.05 (2.09 – 2.05)* | 182.098 – 3.452 (3.692 – 3.452)* |
| Ellipsoidal resolution (Å) | N/A | 3.395 (a*) |
| (direction) |  | 3.395 (b*) |
|  |  | 5.793 (c*) |
| *R*<sub>meas</sub> | 0.096 (2.038) | 0.247 (1.500) |
| *R*<sub>pim</sub> | 0.034 (0.720) | 0.072 (0.561) |
| *I* / $\sigma$(*I*) | 19.5 (1.6) | 8.0 (1.5) |
| *CC*<sub>1/2</sub> | 1.000 (0.628) | 0.998 (0.386) |
| Completeness (spherical) (%) | 100.0 (100.0) | 57.7 (16.1) |
| Completeness (ellipsoidal) (%) | N/A | 93.1 (69.2) |
| Redundancy | 15.0 (15.5) | 12.2 (7.1) |
| **Refinement** |  |  |
| Resolution (Å) | 47.01 – 2.05 | 118.6 – 3.452 |
| No. reflections | 86291 | 22652 |
| *R*<sub>work</sub> / *R*<sub>free</sub> | 0.1942/0.2191 | 0.2633/0.3065 |
| No. atoms | 8118 | 15352 |
| Protein | 7736 | 15352 |
| Ligand/ion | 89 | 0 |
| Water | 293 | 0 |
| *B*-factors | 58.74 | 94.23 |
| Protein | 58.51 | 94.23 |
| Ligand/ion | 90.00 | N/A |
| Water | 55.27 | N/A |
| R.m.s. deviations |  |  |
| Bond lengths (Å) | 0.002 | 0.002 |
| Bond angles (°) | 0.478 | 0.480 |

\*Values in parentheses are for highest-resolution shell.

The electron density maps of the bound BRCA2 Ex14 peptide could not be interpreted unambiguously (Supplementary Figure 1). Nevertheless, we were able to build the BRCA2 Ex14 structure guided by an *Alphafold2* model of the same BRCA2 Ex14 sequence bound to a DMC1 ΔN dimer (Supplementary Figure 2a-e). This resulted in a refined model of the peptide at seven out of the eight DMC1 protomers, with more extensive structural information at instances away from rather than at crystal lattice contacts (Table 1 and Figure 1b). The observed BRCA2 Ex14 peptide contains the PhePP motif (amino-acids 2406-2409), in which F2406 is buried in a hydrophobic pocket and the two consecutive proline residues enable an unusual backbone conformation (Figure 1c). The PhePP-binding site of DMC1 is formed from by two loops, corresponding to amino-acids 147-153 and 178-183. Importantly, the first loop corresponds to one of the most divergent regions between DMC1 and RAD51 (Supplementary Figure 3), which may explain the reported differences in Ex14- and Ex27-binding of RAD51 and DMC1 (Sharan et al., 1997, Esashi et al., 2005, Thorslund et al., 2007). The second loop incudes amino-acid residues V179 (DMC1) and L180 (RAD51) that substantially contribute to the hydrophobic pocket of the differential PhePP-binding site (Figure 1d).

### BRCA2’s PhePP motifs mediate selective RAD51- and DMC1-binding

We next validated the BRCA2-DMC1 structure through interaction studies. Microscale thermophoresis (MST) confirmed that DMC1 ΔN binds to Ex14 (amino-acids 2387-2420) with a *K_D_* of approximately 30 μM, which is over threefold higher affinity than Ex27 (Figure 1e and Supplementary Figure 4). Further, the binding affinity was largely retained for the truncated Ex14-Tr peptide used in crystallographic studies (Figure 1e and Supplementary Figure 4). In contrast, the interaction was largely disrupted upon introduction of alanine mutations of the PhePP motif (herein referred to as Ex14-AAA; amino-acids F2406A, P2408A, P2409A) (Figure 1e). These MST results agree with biochemical pull-down assays, in which DMC1 ΔN was shown to bind to Ex14, Ex14-Tr and BRC4, but not Ex14-AAA or Ex27 (Figure 1f).

To confirm the interaction site on the DMC1 recombinase, we used a mutant in which the loops that form the PhePP-binding site of DMC1 were replaced by the sequences present in RAD51 (herein referred to as DMC1 loop mutant, in which loops 147-GAGGYPG-153 and 178-NVDHDA-183 were mutated to IDRGGGE and GLSGSD, respectively). The DMC1 loop mutant abolished Ex14-binding, confirming that we had identified the correct PhePP-binding site (Figures 1e,g). However, it also failed to interact with Ex27, indicating that additional contacts must contribute to Ex27-binding by RAD51 (Figure 1g). Finally, we used a monomeric DMC1 F85E mutant to demonstrate that Ex14 only binds to oligomeric DMC1, whereas BRC4 binds to both monomeric and oligomeric forms (Figure 1h). Hence, similar to the BRCA2 Ex27-RAD51 interaction (Esashi et al., 2007, Davies and Pellegrini, 2007), we find that BRCA2 Ex14 selectively binds to oligomeric DMC1 at a site formed by residues including V179.

### BRCA2 Ex14 stabilises DMC1 nucleoprotein filaments

What is the function of the BRCA2 Ex14-DMC1 interaction? We used electrophoretic mobility shift assays (EMSAs) to assay the impact of BRCA2-binding on DMC1 nucleoprotein filaments. Firstly, using unfavourable nucleoprotein filament assembly conditions (tris, acetate, EDTA; pH 7.5), we showed that BRCA2 Ex14 promotes the formation of DMC1-ssDNA nucleoprotein filaments (Figure 2a). We then used favourable nucleoprotein filament assembly conditions (triethanolamine + KCl; pH 7.5) to show that BRCA2 Ex14 binds to pre-formed DMC1-ssDNA filaments, inducing a substantial super-shift (Figure 2b). Finally, we demonstrated that binding of BRCA2 Ex14 protects DMC1-ssDNA nucleoprotein filaments from BRC4-mediated disruption (Figure 2c). Importantly, promotion of assembly, stabilisation and protection of DMC1-ssDNA nucleoprotein filaments were abolished by the Ex14-AAA PhePP mutation (Figure 2a-c), indicating that these functions depend on the interaction observed in our BRCA2-DMC1 crystal structure. Hence, BRCA2 Ex14 binds to DMC1 via the PhePP motif, promoting the formation of DMC1-ssDNA nucleoprotein filaments, and protecting filaments from BRC4-mediated disruption, in the same way as was previously observed for the BRCA2 Ex27-RAD51 interaction (Esashi et al., 2007, Davies and Pellegrini, 2007).

**Figure 2.**
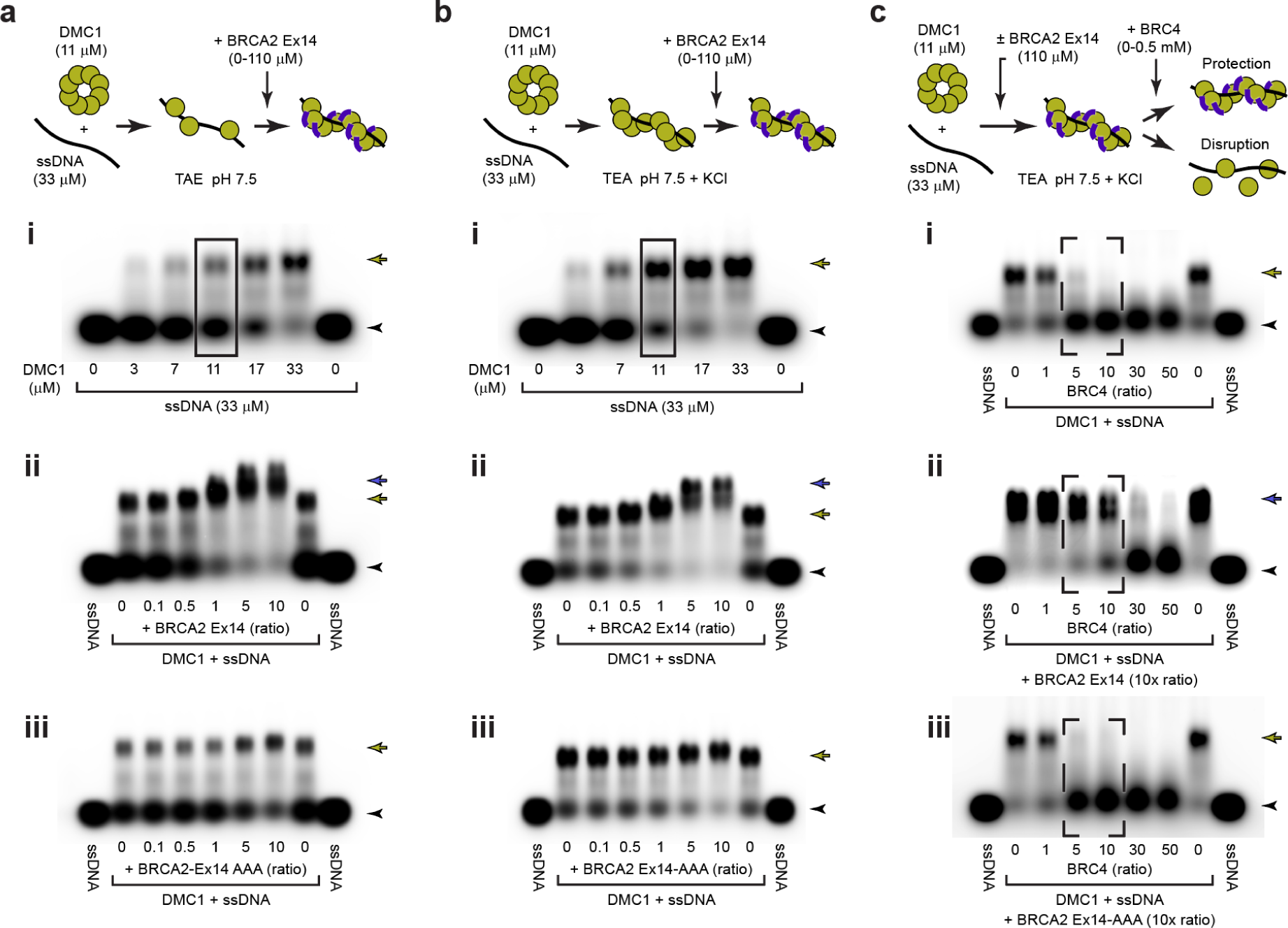
BRCA2 Ex14 stabilises DMC1-ssDNA complexes. Electrophoretic mobility shift assay (EMSAs) analysing the ability of BRCA2 Ex14 to bind, promote and protect DMC1-ssDNA filaments. (**a**) EMSAs using TAE pH 7.5 conditions in which (**i**) DMC1-ssDNA binding is not complete (boxed), demonstrating that (**ii**) BRCA2 Ex14 but not (**iii**) BRCA2 Ex14-AAA mutant promotes the formation of DMC1-ssDNA filaments, using the boxed conditions from panel (**i**). (**b**) EMSAs using TEA pH 7.5 + KCl conditions in which (**i**) DMC1-ssDNA binding is largely complete (boxed), demonstrating that (**ii**) BRCA2 Ex14 but not (**iii**) BRCA2 Ex14-AAA mutant induces a super-shift, demonstrating binding to DMC1-ssDNA filaments, using the boxed conditions from panel (**i**). (**c**) EMSAs using TEA pH 7.5 + KCl conditions in which (**i**) DMC1-ssDNA binding is disrupted by a stoichiometric excess of BRC4, demonstrating that (**ii**) BRCA2 Ex14 but not (**iii**) BRCA2 Ex14-AAA mutant protects against BRC4-mediated disruption (dashed, boxed). BRCA2 peptide concentrations are shown as molar ratios with respect to DMC1 protomers. Arrowheads, free ssDNA; yellow arrows, DMC1-ssDNA complexes; blue arrows, BRCA2-DMC1-ssDNA complexes.

### BRCA2 Ex27 binds to RAD51 through a conserved interaction mode

We wondered whether BRCA2 PhePP motifs bind to RAD51 and DMC1 recombinases through the same interaction mode. Firstly, we built an *Alphafold2* model of BRCA2 Ex27 bound to a RAD51 dimer (Supplementary Figure 5a-e). We then built a model of BRCA2 Ex27 bound to a RAD51 filament by docking our BRCA2-RAD51 model onto a cryo-EM structure of the RAD51 filament (PDB accession 8BSC; Appleby et al., 2023a). We noticed that the N-terminal end of the modelled BRCA2 Ex27 peptide clashed with RAD51 self-association interactions, so removed this and restricted the extent of the Ex27 peptide to amino-acids 3278-3309 (Figure 3a). The resultant model predicts that BRC2 Ex27 binds via precisely the same core PhePP interface as observed in our BRCA2-DMC1 crystal structure (Figure 3b,c). Further, the model predicts that the N-terminal end of BRCA2 Ex27 binds across RAD51’s self-association interface, explaining how it protects filaments from BRC4-mediated disruption (Esashi et al., 2007, Davies and Pellegrini, 2007). Hence, the N-terminal end of BRCA2 Ex14 likely binds across the DMC1 self-association interface in a similar manner to protect from BRC repeat disruption.

**Figure 3.**
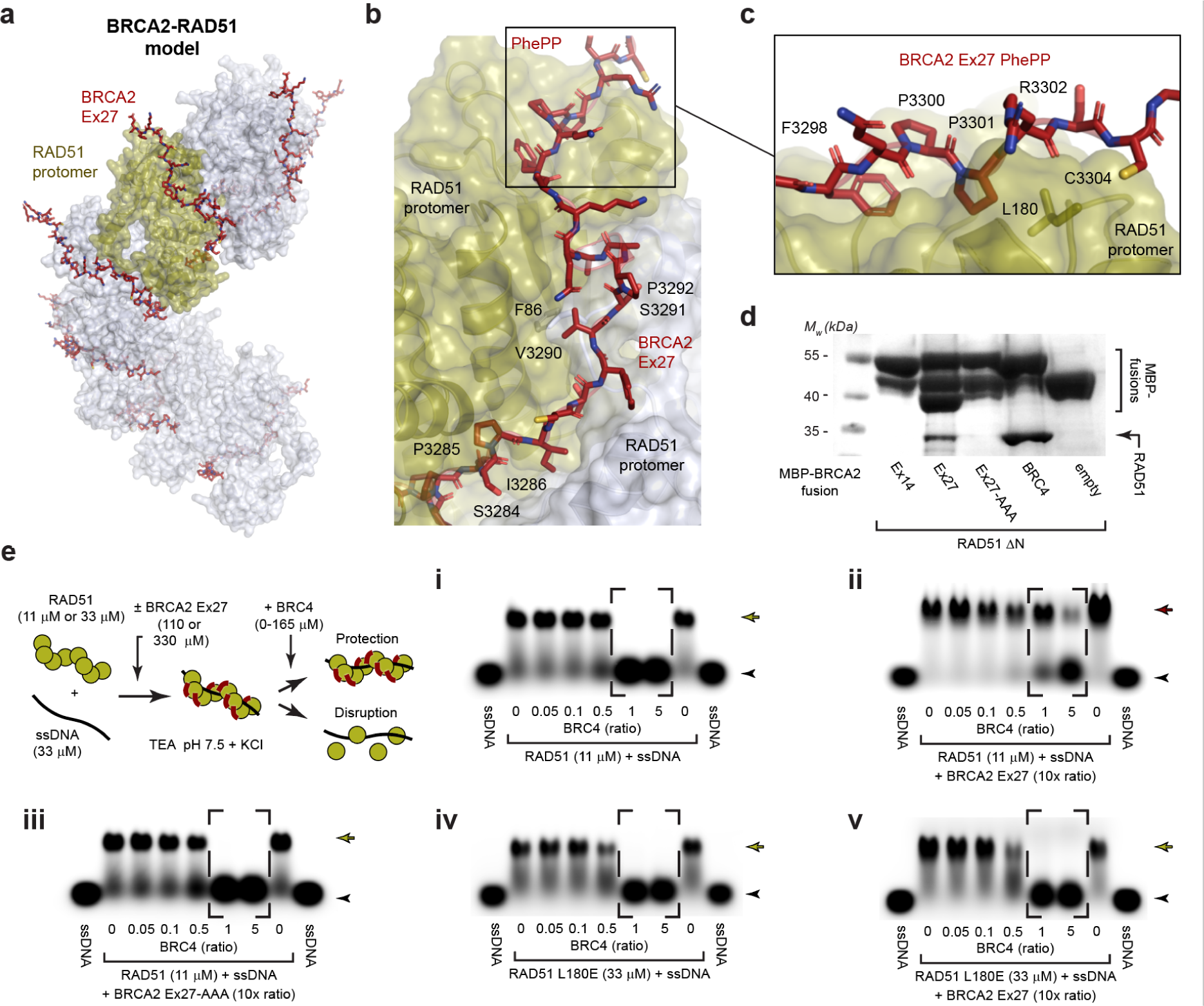
BRCA2 Ex27 stabilises RAD51-ssDNA complexes. **(a)** Model of the BRCA2 Ex27-RAD51 filament structure generated by docking BRCA2 Ex27-RAD51 1:2 complex models onto a previously reported structure of the RAD51 filament (Appleby et al., 2023a; pdb accession: 8BSC). (**b**) The modelled BRCA2 Ex27 peptide includes the PhePP motif (boxed), and a long extension that runs along the interface between adjacent RAD51 promoters, shrouding the F86 self-association interaction. (**c**) The PhePP motif is predicted to bind in the same manner as the BRCA2 Ex14-DMC1 interaction, involving FxPP amino-acids F3298, P3300 and P3301. (**d**) Amylose pull-downs of RAD51 ΔN following recombinant co-expression with MBP-BRCA2 Ex14, Ex27 and its AAA mutant (F3298A, P3300A, P3301A), BRC4 and free MBP (empty). (**e**) EMSAs in which (**i**) RAD51-ssDNA binding is disrupted by a equimolar quantities of BRC4, demonstrating that (**ii**) BRCA2 Ex27 but not (**iii**) BRCA2 Ex14-AAA mutant protects against BRC4-mediated disruption (dashed, boxed), and that (**iv**) RAD51 loop mutant L180E undergoes similar disruption, but (**v**) is not protected by BRCA2 Ex27. BRCA2 peptide concentrations are shown as molar ratios with respect to RAD51 protomers. Arrowheads, free ssDNA; yellow arrows, DMC1-ssDNA complexes; red arrows, BRCA2-RAD51-ssDNA complexes.

We next validated our BRCA2-RAD51 model. Pull-down assays confirmed that RAD51 binds selectively to BRCA2 Ex27 rather than Ex14, in a manner that is disrupted by mutation of the PhePP motif (herein referred to as BRCA2 Ex27-AAA; amino-acids F3308A, P3310A, P3311A) (Figure 3d). Further, we found that protection of RAD51-ssDNA nucleoprotein filaments from BRC4-mediated disruption is abrogated by the BRCA2 Ex27-AAA mutant and by an L180E mutation that targets the divergent loop of RAD51’s PhePP-binding site (Figure 3e). Hence, BRCA2 Ex27 confers protection to RAD51-ssDNA nucleoprotein filaments by binding to RAD51’s PhePP-binding site in the same manner as observed in our crystal structure of BRCA2-DMC1. Thus, we conclude that BRCA2’s PhePP motifs bind selectively to RAD51 and DMC1 nucleoprotein filaments through a conserved interaction mode that stabilises filaments from BRC4-mediated disruption.

## Discussion

The interaction mode between BRCA2’s PhePP motifs and recombinases had remained unknown. Here, we have presented a crystal structure of the BRCA2-DMC1 complex, and solution data, in which we reveal a novel interaction mode, which is conserved between BRCA2’s Ex14 and Ex27 PhePP motifs and recombinases DMC1 and RAD51, respectively. Further, similar PhePP motifs have been reported in other recombinases-interacting proteins, including RAD51AP1 and FIGNL1 (Dunlop et al., 2011, Yuan and Chen, 2013), and are observable in the sequences of other proteins such as SYCP1 (Crichton et al., 2023) and RECQL5 (Islam et al., 2012). Hence, this may be a general mechanism whereby regulatory factors associate with RAD51 and DMC1 nucleoprotein filaments. Overall, our data support a dichotomy, in which BRCA2’s BRC and PhePP motifs have distinct roles in remodelling recombinases and stabilising nucleoprotein filaments, respectively, in both mitotic and meiotic recombination.

How does BRCA2 PhePP-binding confer stability to RAD51 and DMC1 nucleoprotein filaments? The BRCA2-DMC1 crystal structure reported herein shows only the core PhePP interface, which is distant from the recombinase self-association interface. Nevertheless, the BRCA2 Ex27-RAD51 model suggests that the upstream sequence binds across the self-association interface, which would sterically hinder access to the self-association interface, thereby explaining its stabilisation and protection from BRC-mediated disruption (Esashi et al., 2007, Davies and Pellegrini, 2007). In a parallel study, a cryo-EM structure of the BRCA2 Ex27-RAD51 complex was reported (Appleby et al., 2023b), which agrees with the model presented herein. Further, in the cryo-EM structure and in our model, the positively charged N-terminal end of Ex27 is located close to the filament axis, where bound DNA is located. Hence, we suggest that Ex27 further stabilises RAD51 nucleoprotein filaments by its N-terminus binding directly to DNA, in keeping with recent biochemical findings (Kwon et al., 2023). It is likely that BRCA2 Ex14 confers stability and protection to DMC1 nucleoprotein filaments in a similar manner to Ex27, through its upstream sequence binding across the self-association interface.

There are some notable differences in the sequences of BRCA2 Ex14 and Ex27, which must affect their function. Firstly, Ex14 show little conservation outside the PhePP motif (Thorslund et al., 2007), suggesting that whilst it must extend across the self-association interface to confer protection, it forms fewer recombinase interactions than Ex27. Secondly, Ex14 lacks the CDK phosphorylation site that is present in Ex27 and is responsible for disrupting binding at the G2-M transition (Esashi et al., 2005). This likely represents the difference between the BRCA2 Ex14-DMC1 interaction functioning in meiosis, and the BRCA2 Ex27-RAD51 interaction functioning in replication fork preservation (Feng and Jasin, 2017, Tye et al., 2021, Kwon et al., 2023).

Whilst BRCA2 PhePP interactions have thus far been considered in isolation, they likely operate as part of a wider ‘recombinosome’ structure. BRCA2’s PhePP motifs are located on either side of its C-terminal DNA-binding domain (Yang et al., 2002). Hence, filament-stabilising interactions by Ex14 and Ex27 may occur cooperatively with DNA-binding and nucleation of nucleoprotein filament formation by BRCA2, which is known to involve displacement of ssDNA-binding protein RPA (Shahid et al., 2014, Bell et al., 2023). Further, Ex14 is located adjacent to a binding site of meiotic recombination complex MEILB2-BRME1 (Zhang et al., 2019, Zhang et al., 2020), which is proposed to displace meiosis-specific ssDNA-binding complex MEIOB-SPATA22 and act as DNA clamp (Zhang et al., 2022, Gurusaran et al., 2023). Hence, BRCA2 may coordinate the formation of a ‘meiotic recombinosome’ in which the key functionalities of meiotic recombination are brought together to enable the loading and stabilisation of DMC1 nucleoprotein filaments for inter-homologue recombination. Our understanding of meiotic recombination would be truly transformed by structure elucidation of this multi-protein assembly.

## Materials and Methods

### Recombinant protein expression and purification

The sequences corresponding to human DMC1 (amino acids 1-340) and DMC1 ΔN (83-340) were cloned into pHAT4 (Peranen et al., 1996) for expression as a TEV-cleavable His6-tag fusion in BL21 (DE3) cells (Novagen®) in 2xYT media, induced with 0.5 mM IPTG for 16 hours at 25°C. Cells were lysed by sonication in 20 mM Tris pH 8.0, 500 mM KCl and fusion proteins were purified from clarified lysate through consecutive Ni-NTA (Qiagen) and HiTrap Q HP (GE Healthcare) ion exchange chromatography. His6-tag removal was mediated by incubation overnight with TEV protease. Cleaved DMC1 was further purified by HiTrapQ HP ion exchange chromatography and size exclusion chromatography (HiLoad^TM^ 16/600 Superdex 200, GE Healthcare) in 20 mM Tris pH 8.0, 150 mM KCl, 2 mM DTT.

RAD51 (amino acids 1-339) and RAD51 L180E were co-expressed with an MBP-fusion of BRCA2 BRC4 (amino acids 1519-1551) in BL21 (DE3) cells (Novagen®) in 2xYT media, and induced with 0.5 mM IPTG for 16 hours at 25°C. Cells were lysed by sonication in 20 mM Tris pH 8.0, 500 mM KCl, and the complex of RAD51 and MBP-BRCA2 BRC4 was purified from clarified lysate through consecutive amylose and Ni-NTA (Qiagen) affinity chromatography. The complex was disrupted through HiTrap Q HP (GE Healthcare) ion exchange chromatography, and RAD51 was further purified through HiTrap Heparin HP (GE Healthcare) ion-exchange chromatography and size exclusion chromatography (HiLoad^TM^ 16/600 Superdex 200, GE Healthcare) in 20 mM Tris pH 8.0, 150 mM KCl, 2 mM DTT.

The sequences corresponding to human BRCA2 Ex14 (amino acids 2387-2420) and BRCA2 Ex14-AAA were cloned into pMAT11 for expression as TEV-cleavable His6-MBP-tag fusions in BL21 (DE3) cells (Novagen®) in 2xYT media, induced with 0.5 mM IPTG for 16 hours at 25°C. Cells were lysed by sonication in 20 mM Tris pH 8.0, 500 mM KCl and the fusion proteins purified from clarified lysate through consecutive Ni-NTA (Qiagen) and HiTrap Q HP (GE Healthcare) ion exchange chromatography. His6-MBP-tag removal was mediated by incubation overnight with TEV protease. Cleaved BRCA2 Ex14 was further purified by HiTrapSP HP ion exchange chromatography and size exclusion chromatography (HiLoad^TM^ 16/600 Superdex 200, GE Healthcare) in 20 mM Tris pH 8.0, 150 mM KCl, 2 mM DTT.

All other BRCA2 peptide sequences utilised in experiments (other than affinity pulldown experiments) were synthesized by Severn Biotech including BRC4 (amino acids 1519-1551), BRCA2 Ex14-Tr (amino acids 2401-2414), BRCA2 Ex27 (3270-3315) and BRCA2 Ex27-AAA.

Protein samples were concentrated using 10,000/3,000 MWCO centrifugal units (Amicon) and were stored at -80<C following flash-freezing in liquid nitrogen. Protein samples were analysed by SDS-PAGE with Coomassie staining, and concentrations were determined by UV spectroscopy using a Cary 60 UV spectrophotometer (Agilent) with extinction coefficients and molecular weights calculated by ProtParam (http://web.expasy.org/protparam/).

For affinity pulldown experiments, MBP-fusions of BRCA2 fragments (Ex14, amino-acids 2387-2420, Ex14-Tr, amino acids 2401-2414, Ex27, amino acids 3270-3315, BRC4, amino acids 1519-1551) were cloned into pMAT11 vectors (Peranen et al., 1996) and co-expressed with DMC1 core, DMC1 core loop mutant (in which DMC1 amino acids 147-GAGGYPG-153 and 178-NVDHDA-183 were replaced by the cognate sequences of RAD51 148-IDRGGGE-154 and 179-GLSGSD-184), RAD51 core, RAD51 core loop mutant (the reverse switch of DMC1 core loop mutant), and RAD51. MBP-fusions and complexes were purified through amylose affinity chromatography.

### Crystallisation and structure solution of DMC1 (6R3P)

DMC1 ΔN protein crystals were obtained through vapour diffusion in hanging drops, by mixing 1 µl of protein at 15 mg/ml with 1 µl of crystallisation solution (50 mM HEPES-NaOH pH 7.2, 50 mM MgCl_2_, 500 mM NaCl, 7.5 % PEG3350) and equilibrating at 20°C for 2 weeks. Crystals were cryo-protected using 6 M sodium formate and cryo-cooled in liquid nitrogen. X-ray diffraction data were collected at 0.9159 Å, 100 K, as 2000 consecutive 0.10° frames of 0.010 s exposure on a Pilatus 6M detector at beamline I04-1 of the Diamond Light Source synchrotron facility (Oxfordshire, UK). Data were indexed, integrated in XDS (Kabsch, 2010), scaled in XSCALE (Diederichs et al., 2003), and merged in Aimless (Evans, 2011). Crystals belong to orthorhombic spacegroup I422 (cell dimensions a = 174.49 Å, b = 174.49 Å, c = 179.57 Å, α = 90°, β = 90°, γ = 90°), with four DMC1 protomers in the asymmetric unit. Structure solution was achieved through molecular replacement using PHASER (McCoy et al., 2007), with a single chain from pdb accession 4HYY as a search model. Model building was performed through iterative re-building by PHENIX Autobuild (Adams et al., 2010) and manual building in Coot (Emsley et al., 2010), with the addition of PEG ligands. The structure was refined using PHENIX refine (Adams et al., 2010), using isotropic atomic displacement parameters with two TLS groups per chain. The structure was refined against data to 2.05 Å resolution, to R and R_free_ values of 0.1942 and 0.2191 respectively, with 99.07% of residues within favoured regions of the Ramachandran plot (0 outliers), clashscore of 1.28and overall MolProbity score of 0.85 (Chen et al., 2010).

### Crystallisation and structure solution of a BRCA2-DMC1 complex (8R2G)

DMC1 ΔN protein crystals were obtained through vapour diffusion in hanging drops, by mixing 1 µl of protein at 20 mg/ml with 1 µl of crystallisation solution (50 mM HEPES-NaOH pH 7.4, 50 mM MgCl_2_, 500 mM NaCl, 8 % PEG 3350) and equilibrating at 20°C for 2 weeks. Crystals were soaked overnight in crystallisation solution containing 3.5 mM BRCA2 Ex14-Tr peptide (RPTKVFVPPFKTKS; synthesised by Severn Biotech). Crystals were cryo-protected using 20 % glycerol and cryo-cooled in liquid nitrogen. X-ray diffraction data were collected at 0.9795 Å, 100 K, as 2000 consecutive 0.10° frames of 0.050 s exposure on a Pilatus3 6M detector at beamline I04 of the Diamond Light Source synchrotron facility (Oxfordshire, UK). Data were processed using AutoPROC (Vonrhein et al., 2011), in which indexing, integration, scaling and merging were performed by XDS (Kabsch, 2010) and Aimless (Evans, 2011), and anisotropic correction with a local I/σ(I) cut-off of 1.2 was performed by STARANISO (Tickle et al., 2018). Crystals belong to tetragonal spacegroup P4_1_22 (cell dimensions a = 125.40 Å, b = 125.40 Å, c = 364.195 Å, α = 90°, β = 90°, γ = 90°), with one DMC1 octamer in the asymmetric unit. Structure solution was achieved through molecular replacement using Phaser (McCoy et al., 2007), through placement of four DMC1 dimers from its high-resolution structure (pdb accession 6R3P). The intact molecular replacement solution was refined using PHENIX refine (Adams et al., 2010) and alternative side-chain conformations were removed. BRCA2 peptides were built using *Alphafold2* models of a BRCA2-DMC1 1:2 complex, which were docked onto the DMC1 protomers of the octameric ring using *PyMOL.* The completed BRCA2-DMC1 structure was refined using PHENIX refine (Adams et al., 2010), with isotropic atomic displacement parameters, using reference model restraints from the high-resolution DMC1 structure (pdb accession 6R3P). The structure was refined against anisotropy-corrected data with resolution limits between 3.4 Å and 5.8 Å, to R and R_free_ values of 0.2635 and 0.3057 respectively, with 98.80% of residues within the favoured regions of the Ramachandran plot (0 outliers), clashscore of 2.88 and overall MolProbity score of 1.08 (Chen et al., 2010).

### Electrophoretic mobility shift assays (EMSAs)

To form nucleoprotein filaments, DMC1 and RAD51 was incubated with fluorescent 100-base ssDNA (5’-FAM-AATTCTCATTTTACTTACCGGACGCTATTAGCAGTGGGTGAGCAAAAACAGGAAGGCAAAATGCCGCAAAAAAGGGAATAAGGGCGACACGGAAATGTTG-3’) at the concentrations indicated in the figures for 30 minutes at 4°C in 50 mM TEA (triethanolamine) pH 7.5, 1 mM ATP, 2 mM MgCl2, 0.1 mg/ml BSA, 2 mM DTT, 200 mM KCl. For assays requiring incomplete nucleoprotein filament assembly, the reaction buffer consisted of 50 mM TAE (tris, acetate, EDTA) pH 7.5, 2 mM ATP, 2 mM MgCl2, 2 mM DTT. In nucleoprotein formation stimulation assays or supershift experiments, BRCA2Ex14 or BRCA2Ex14-AAA were added for a further 30 min at 4°C. For protection experiments, the DMC1, RAD51, or RAD51 L180E–DNA complexes were incubated with BRCA2 Ex14, Ex14-AAA, Ex27, or Ex27-AAA for 30 min at 4°C prior to the addition of BRC4 and further incubation for 30 min at 4°C (concentrations provided in figures). Glycerol was added at a final concentration of 3%, and samples were analysed by electrophoresis on a 1% (w/v) agarose gel in 0.5x TAE pH 8.0 at 20-40 V for ∼4 h at 4 °C. DNA was detected by FAM using a TyphoonTM FLA 9500 (GE Healthcare), with 473 nm laser at excitation wavelength 490 nm and emission wavelength 520 nm, using the LPB filter and a PMT voltage of 500 V.

### Structural modelling

Models were generated using a local installation of *Alphafold2* (Jumper et al., 2021). Models of BRCA2-DMC1 and BRCA2-RAD51 complexes were generated using a 1:2 ratio of BRCA2 Ex14/Ex27 and DMC1 core and RAD51, respectively. BRCA2-RAD51 models were docked onto a RAD51 filament (Appleby et al., 2023a; pdb accession: 8BSC), and clashing regions at the N-termini (up to amino-acid 3277) of modelled BRCA2 Ex27 peptides were removed. For comparison between PhePP-binding loops, the BRCA2-DMC1 crystal structure was superposed with the RAD51 core (Pellegrini et al., 2002; pdb accession 1N0W) using *PyMOL*.

### Protein sequence and structure analysis

Multiple sequence alignments were generated using *Jalview*(Waterhouse et al., 2009), and molecular structure images were generated using the *PyMOL* Molecular Graphics System, Version 2.0.4 Schrödinger, LLC.

### Accession codes and data availability

Crystallographic structure factors and atomic co-ordinates have been deposited in the Protein Data Bank (PDB) under accession numbers 6R3P and 8R2G.

## Acknowledgements

We thank Diamond Light Source and the staff of beamlines I04 and I04-1 (proposals mx13587 and mx18598). We thank A. Baslé for assistance with X-ray crystallographic and CD data collection. J.M.D. is a Herchel Smith Fellow. O.R.D. is a Wellcome Senior Research Fellow (Grant Number 219413/Z/19/Z).

## Author contributions

J.M.D. performed experimental work. O.R.D. solved the crystal structures and wrote the manuscript. J.M.D. and O.R.D. designed experiments and analysed data.

---

**Supplementary Figure 1.**
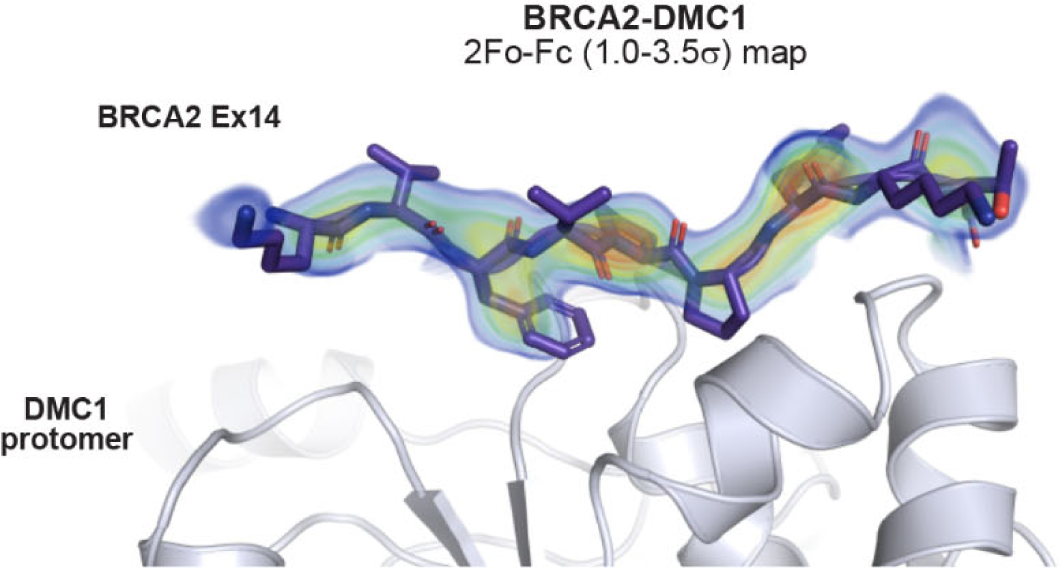
Crystal structure of the BRCA2-DMC1 core complex. 2Fo-Fc electron density map of the BRCA2-DMC1 core structure, presented as a rainbow between 1.0σ (blue) and 3.5σ (red), superimposed on the refined crystallographic model.

**Supplementary Figure 2.**
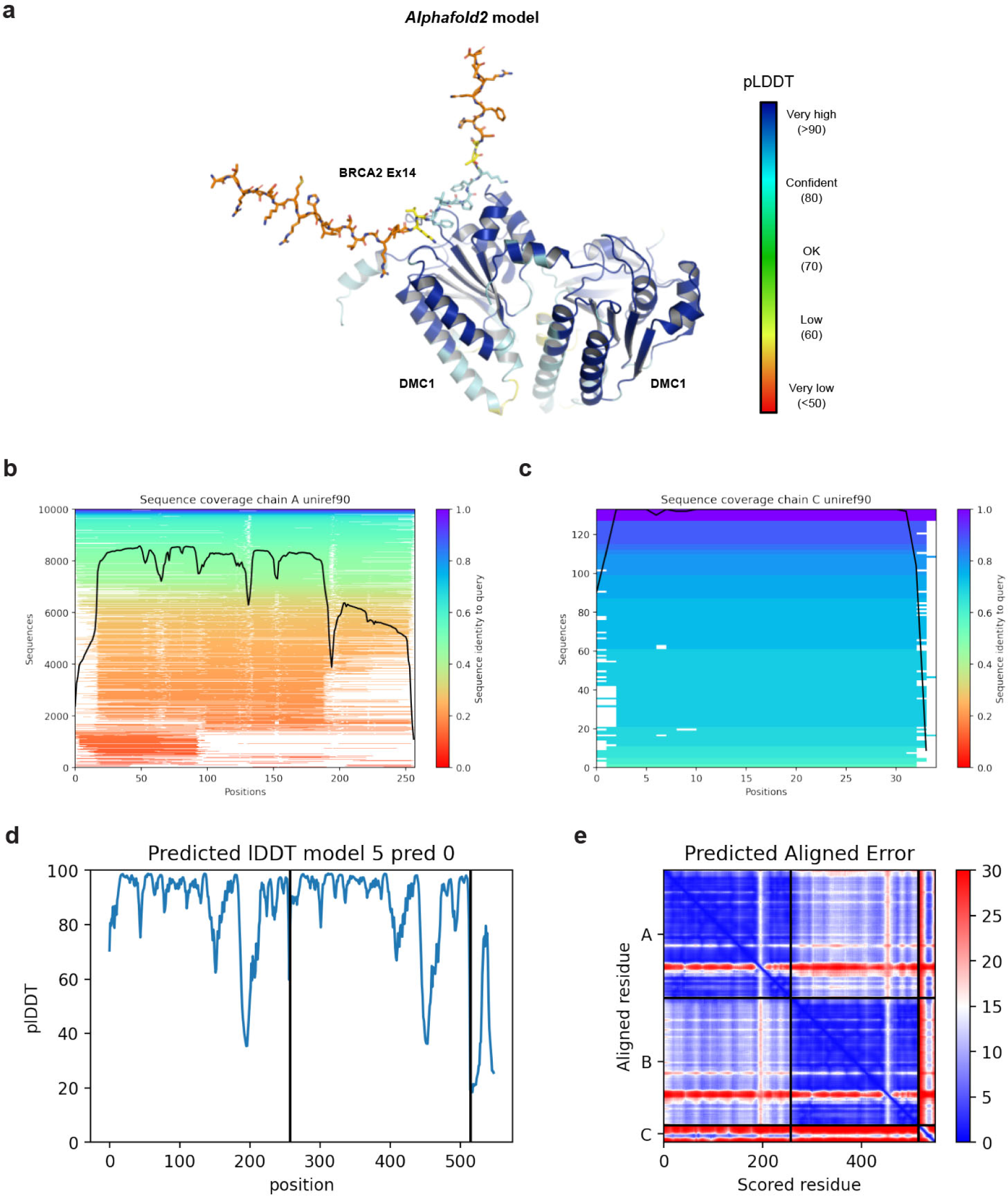
*Alphafold2* model of the BRCA2 Ex14-DMC1 core 1:2 complex. **(a)** *Alphafold2* model of the BRCA2 Ex14-DMC1 core 1:2 complex coloured according to predicted LDDT (pLDDT) scores, between blue (>90) and red (<50). (**b**) Representations of the multiple sequence alignments generated and used by *Alphafold2*, showing the number of sequences and sequence identity against the position along the (**c**) DMC1 core and (**d**) BRCA2 Ex14 query sequences. (**e**) Predicted LDDT (pLDDT) scores shown for each amino-acid of the two DMC1 core and one BRCA2 Ex14 chains. (**f**) Predicted aligned error scores between each amino-acid of the two DMC1 core and one BRCA2 Ex14 chains, between blue (low error) and red (high error).

**Supplementary Figure 3.**
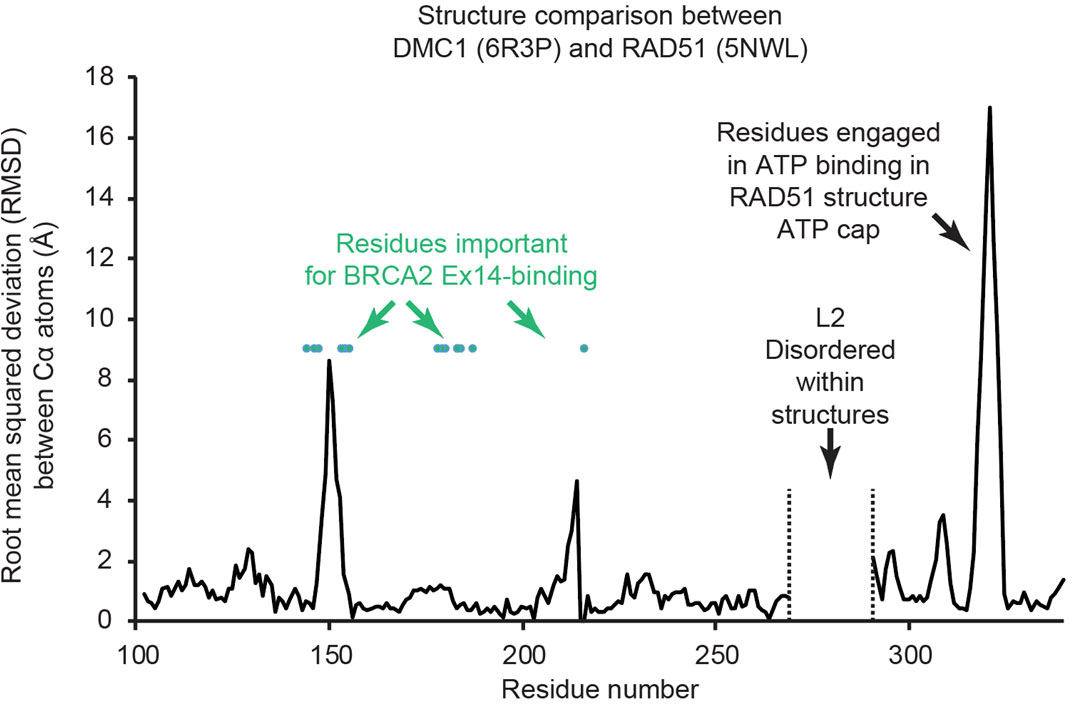
Conservation of sequences within DMC1 and RAD51 recombinases. Root mean squared deviation (RMSD) between Cα atoms of DMC1 and RAD51, using structures 6R3P (this study) and 5NWL (Brouwer et al., 2018).

**Supplementary Figure 4.**
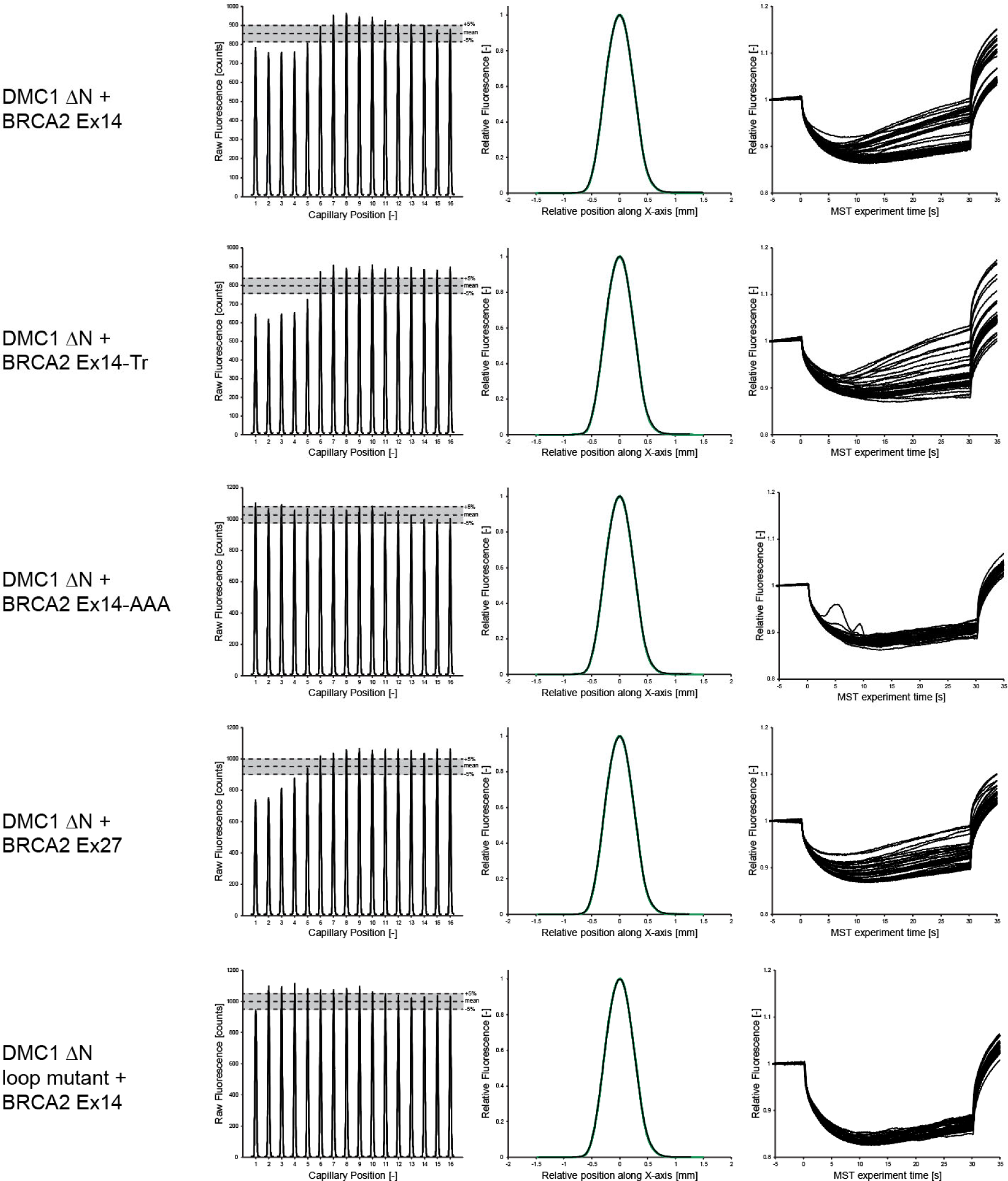
Microscale thermophoresis (MST) analysis of the BRCA2 Ex14-DMC1 interaction. MST analysis corresponding to Figure 1e. Initial fluorescence (left), individual data series (middle) and relative fluorescence for each of the three data series (right), for each BRCA2 peptide and DMC1 ΔN construct, as indicated.

**Supplementary Figure 5.**
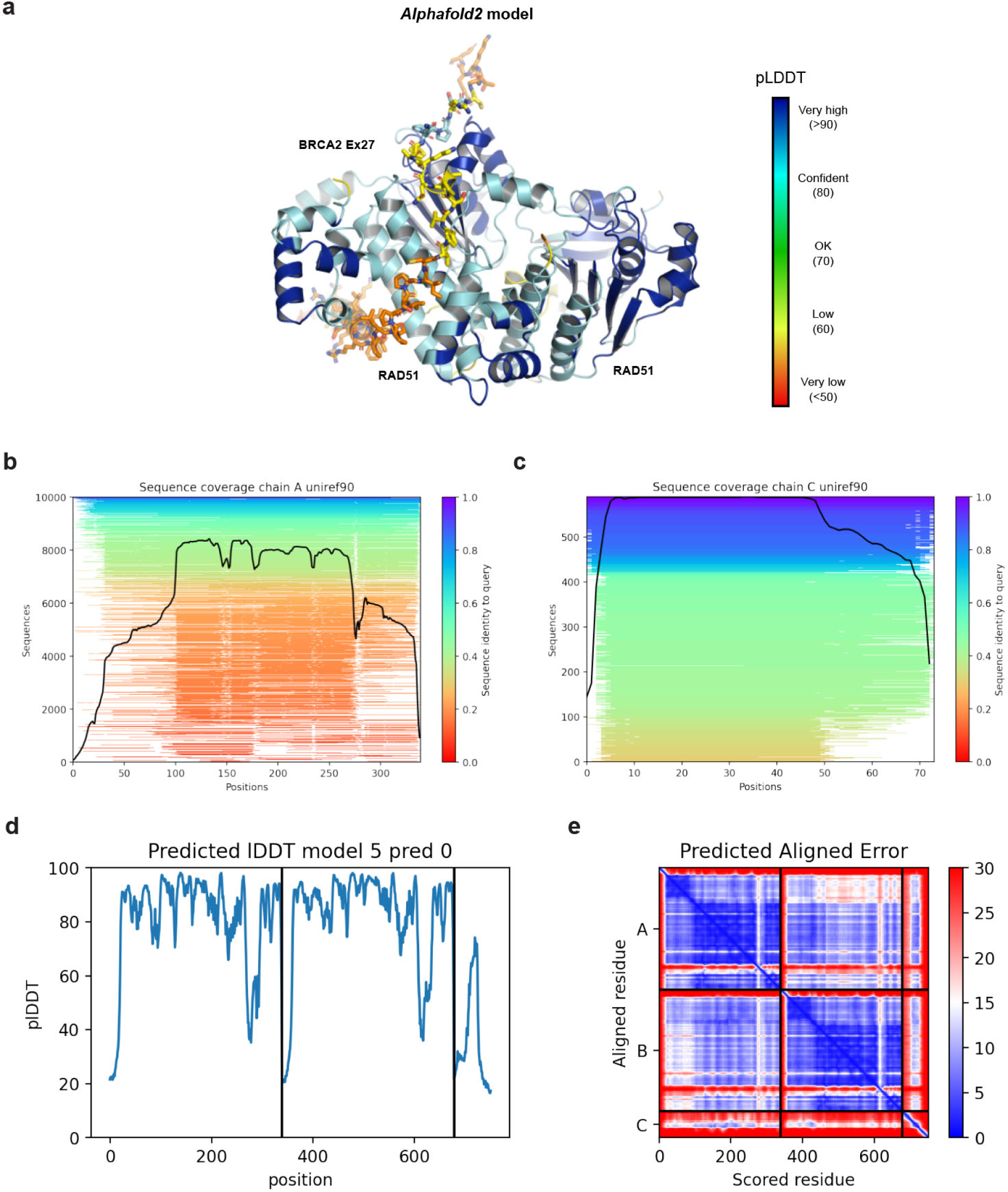
*Alphafold2* model of the BRCA2 Ex27-RAD51 core 1:2 complex. **(a)** *Alphafold2* model of the BRCA2 Ex27-RAD51 core 1:2 complex coloured according to predicted LDDT (pLDDT) scores, between blue (>90) and red (<50). (**b**) Representations of the multiple sequence alignments generated and used by *Alphafold2*, showing the number of sequences and sequence identity against the position along the (**c**) RAD51 and (**d**) BRCA2 Ex27 query sequences. (**e**) Predicted LDDT (pLDDT) scores shown for each amino-acid of the two RAD51 and one BRCA2 Ex27 chains. (**f**) Predicted aligned error scores between each amino-acid of the two RAD51 and one BRCA2 Ex27 chains, between blue (low error) and red (high error).

